# Bringing BOS to light: Uncovering the key enzyme in the biosynthesis of the neurotoxin β-ODAP in Grass Pea (*Lathyrus sativus* L.)

**DOI:** 10.1101/2020.11.29.402396

**Authors:** Moshe Goldsmith, Shiri Barad, Maor Knafo, Alon Savidor, Shifra Ben-Dor, Alexander Brandis, Tevie Mehlman, Yoav Peleg, Shira Albeck, Orly Dym, Efrat Ben-Zeev, Ziv Reich

**Author notes:** **Corresponding authors:** Moshe Goldsmith and Ziv Reich, Department of Biomolecular Sciences, Weizmann Institute of Science, Rehovot 7610001, Israel. These authors contributed equally to this work (Moshe Goldsmith, Shiri Barad, Maor Knafo). **Summary:** The identification and characterization of a long-sought enzyme from grass pea (*Lathyrus sativus* L.).

## Abstract

Grass pea (*Lathyrus sativus* L.) is a grain legume commonly grown in parts of Asia and Africa for food and forage. While being a highly nutritious and robust crop, able to survive both drought and floods, it produces a neurotoxic compound, β-*N*-oxalyl-L-α,β-diaminopropionic acid (β-ODAP), which can cause a severe neurological disorder if consumed as a main diet component. So far, the enzyme that catalyzes the formation of β-ODAP has not been identified. By combining protein purification and enzymatic assays with transcriptomic and proteomic analyses, we were able to identify the enzyme β-ODAP synthetase (BOS) from grass pea. We show that BOS is an HXXXD-type acyltransferase of the BAHD superfamily and that its crystal structure is highly similar to that of plant hydroxycinnamoyl transferases. The identification of BOS, more than 50 years after it was proposed, paves the way towards the generation of non-toxic grass pea cultivars safe for human and animal consumption.

## Report

Grass pea (*Lathyrus sativus* L.) is an annual legume crop grown for food and forage, mostly in South Asia and Sub-Saharan Africa (*1, 2*). Its attractiveness as a farming crop stems from its tolerance to harsh environmental conditions such as drought, high salinity and flooding (*3–5*), its resistance to insects and fungal diseases (*6, 7*), as well as its high grain-yield (*8*) and high nutritional value (*9, 10*). Unfortunately, it also produces a neurotoxic glutamate analogue, named β-*N*-oxalyl-L-α,β-diaminopropionic acid (β-ODAP) (*11, 12*). This neurotoxin may cause lathyrism, a neurodegenerative disorder characterized by lower limb paralysis (*13, 14*), if consumed as a primary diet component over a prolonged period of time. Breeding attempts have been successful at generating grass pea cultivars with reduced concentrations of β-ODAP (*15*). However, none of the cultivars generated so far are completely devoid of β-ODAP and toxin levels may increase under stress conditions, such as drought (*16*). A constitutively expressed fungal oxalate decarboxylase gene, introduced into grass pea to degrade oxalate and, subsequently, reduce the concentration of oxalyl-CoA, a precursor of β-ODAP (*17*), had likewise failed to eliminate β-ODAP production. Grass pea, therefore, remains an underutilized crop of limited economic importance in global markets, yet an often-essential source of food and income security for resource-poor farmers in developing countries (*2, 18*).

The biological role of β-ODAP in grass pea is still unknown. It has been proposed to transport zinc in soils depleted of zinc or rich in iron (*19*), to act as a radical scavenger or to enhance symbiosis with *Rhizobium* bacteria (*16, 20, 21*). The biosynthesis of β-ODAP in grass pea was studied more than 50 years ago (*22*). It was suggested to occur through the ligation of oxalyl-CoA and L-α,β-diaminopropionic acid (L-DAPA), catalyzed by a putative β-ODAP synthetase (**Fig 1**) (*23*). However, this synthetase has hitherto not been identified or characterized.

**Figure 1.**
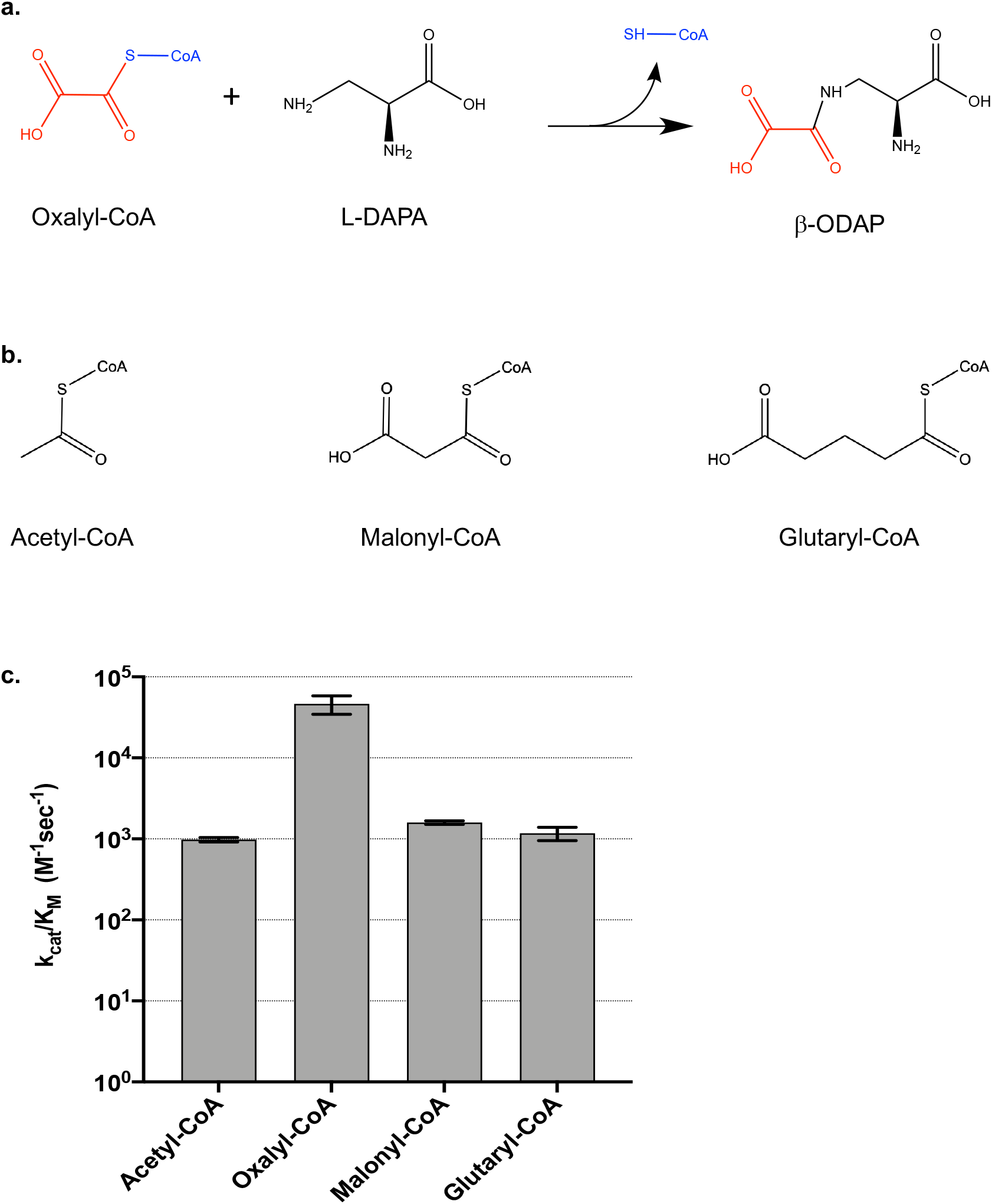
Reaction scheme and catalytic efficiencies of BOS. **a.** A scheme of the reaction catalyzed by BOS. **b.** The structures of additional CoA-substrates studied in this work. **c.** Catalytic efficiency values of BOS with different CoA substrates. The values were determined at varying concentrations of L-DAPA and saturating concentrations of the CoA substrates and are, therefore, considered apparent.

Here, we used grass pea seeds and seedlings in order to identify and isolate β-ODAP synthetase (BOS) by fractionation and protein purification. To trace the enzyme during purification, we assayed protein fractions for their ability to ligate oxalyl-CoA to L-DAPA. Oxalyl-CoA was synthesized *in-vitro* from oxalic acid and CoA, using an oxalyl CoA-synthetase that was also purified from grass pea (*Ls*OCS) (*24*). Following chromatographic separations on multiple columns (**Materials and Methods**), we obtained four enzymatically active fractions. These fractions were subjected to a proteomic LC-MS/MS analysis. Since the genomic sequence of grass pea was unknown at the time, we sequenced grass pea mRNA using long-read PacBio sequencing. The constructed set of annotated full-length mRNA transcripts was translated and used as a database for the identification of grass pea proteins in the enzymatically active fractions (**Table S1**). Between 247 and 1969 different proteins, including BOS, were identified in the different active fractions. However, only 222 proteins were found in all four fractions purified from grass pea (**Table S2**). Of these, 40 protein sequences were predicted to be ligases, transferases, synthetases or CoA-binding enzymes. We selected 14 sequences of CoA-binding and Shikimate pathway ligases and transferases that seemed likely to bind the substrates and catalyze the ligation reaction for functional analysis. Selected genes were synthesized, cloned and expressed in *E.coli* cells and their levels of BOS activity were assayed in clarified cell lysates. Of the 14 genes cloned and tested, only one exhibited significant activity (**Fig. S1**).

To verify that the observed activity of the clone was that of a bona fide β-ODAP synthetase, the recombinant enzyme was purified from *E.coli* cells and its enzymatic activity was assayed *in-vitro*. The enzyme purified as a stable monomer of 439 aa (49.3 kDa) (**Fig. S2**) and exhibited a moderate catalytic efficiency with an apparent turnover rate (*k*_cat_) and Michaelis (*K*_M_) constants of 118 ± 15 [sec^−1^] and 2.54 ± 5.7 [mM], respectively, giving rise to an apparent catalytic efficiency (*k*_cat_/*K*_M_) of 4.6 ± 1.2 ×10^4^ [sec^−1^M^−1^] (**Fig. S3**) and a specific activity of 13.2 ± 1.6 [M·sec^1^·gr^−1^]. When oxalyl-CoA was replaced by similar plant metabolites such as acetyl-CoA, malonyl-CoA or glutaryl-CoA, the catalytic efficiencies dropped by 30 to 48-fold (**Fig. 1, S3**); indicating enzymatic specificity towards oxalyl-CoA, the native substrate. To verify the formation of β-ODAP in the reaction, we examined the concentrations of L-DAPA, β-ODAP and α-ODAP, the non-toxic isomer of β-ODAP, using reversed-phase chromatography and mass spectrometry (LC-MS). The results (**Fig. S4**) showed that >99.9% of the L-DAPA in the reaction mixture had converted to β-ODAP while the concentrations of L-DAPA in the negative control reactions remained unchanged and the concentrations of α-ODAP in all samples were extremely low. Thus, BOS displays a high degree of stereospecificity towards the formation of β-ODAP.

We analyzed the domain sequences of BOS to determine which family it belongs to and to identify putative homologs in other species. Almost the entire length of the protein (residues 3-434 out of 439, full length) comprises a transferase domain (PF02458) of a family found predominantly in plants and fungi. Database searches showed that the closest sequences are related to EPS1 (Enhanced Pseudomonas Susceptibility 1) - an *Arabidopsis thaliana* protein that belongs to the BAHD acyltransferase superfamily (*25*). We found two hallmarks of BAHD superfamily enzymes in BOS: A conserved active site HXXXD motif (residues 162-166, HSVVD, **Fig. 2**) and a structural DFGWG motif (residues 381-385, **Fig. S5**) (*25–27*).

**Figure 2.**
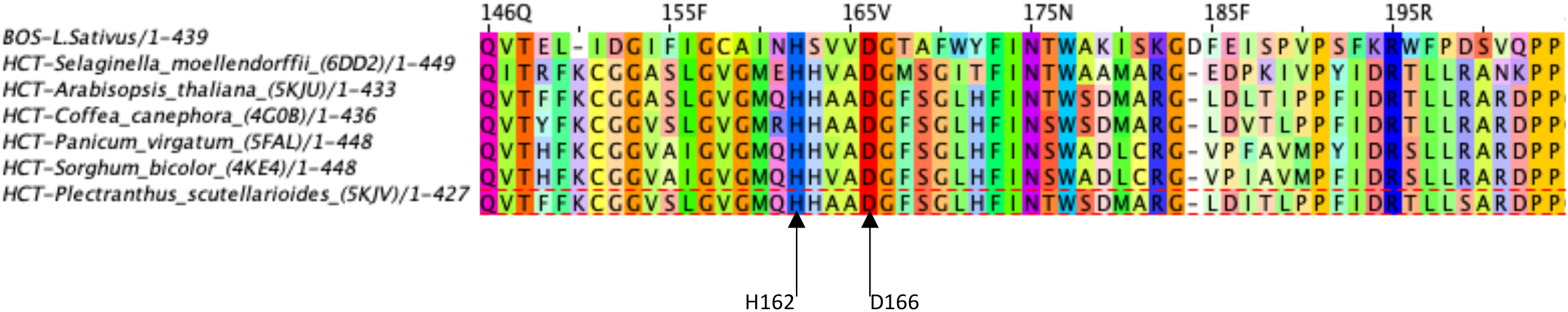
Sequence alignment of the HXXXD motif in BOS and several HCT homologs. The sequences of CoA:shikimate hydroxycinnamoyl transferases (HCTs) from different species were aligned to that of BOS. The black arrows denote His162 and Asp166 in BOS, which are part of the conserved BAHD acyltransferase HXXXD motif. In parentheses are the PDB codes of the proteins.

A search for other BAHD acyltransferases in the grass pea transcriptome identified 30 additional members (**Appendix A**). To identify the clade to which BOS belongs, a phylogenetic tree of BAHD proteins was constructed (**Fig. 3**), using the 31 BAHD proteins from grass pea, 47 BAHD proteins from *Medicago trunculata*, 58 BAHD proteins from *Arabidopsis thaliana*, 96 from *Populus trichocarpa* (*28*), and 67 sequences with known functions, from other species (*25, 29*). In total, 299 sequences were aligned, 93 of which had known biological functions (**Appendix A, B**). The tree (**Fig. 3**) shows the typical clades of the superfamily, namely, Ia, Ib, II, IIIa IIIb, IV, Va and Vb (*25, 28, 29*) with many branches exhibiting species-specific expansions (**Fig. S6**). BOS, along with 10 other grass pea proteins forms part of clade Ib, and is the second protein in that clade, apart from *A. thaliana* DCR (*30*), to be characterized.

**Figure 3.**
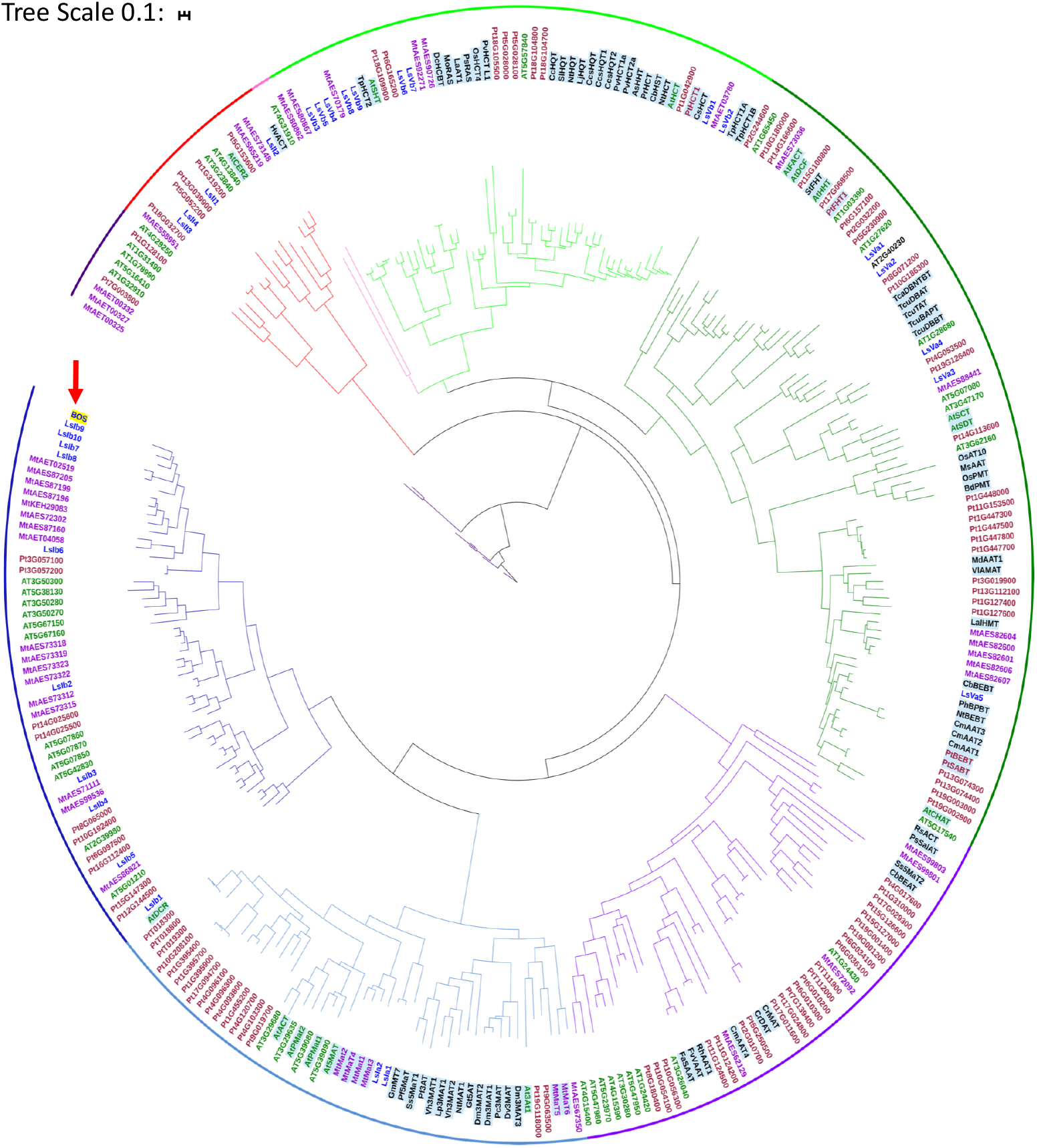
Phylogenetic tree of BAHD superfamily proteins. Sequence names are prefixed by a two-letter species-code and colored by species: *L. sativus* (Ls) blue*, A. thaliana* (At) green, *M. trunculata* (Mt) purple, *P. trichocarpa* (Pt) brown; others species are colored black (full details in **Supplemental Appendix A**). The outer ring and clades are colored as follows (clockwise from the open end on the left): putative clade IIIb (dark purple), clade II (red), clade IV (pink), clade Vb (light green), clade Va (dark green), clade IIIa (light purple), clade Ia (light blue), clade Ib (dark blue). Names of sequences with known function are shaded in light blue; BOS is marked by a red arrow and shaded in yellow.

BOS was crystalized and the structure was solved to 2.35 Å resolution (**Table S3**). The enzyme consists of 15 β-strands and 11 α-helices and is organized into two, equally sized domains of ~200 amino acids. Each domain centers around a 6-stranded β-sheet, flanked by α-helices (**Fig. 4a**). The two domains are connected by two loops, one (residues 183-209) links the N- and C-terminal domains (**Fig. 4a**), and the other (residues 371-388) joins β-strand 13 from the C-terminal domain (residues 389-393) to the β-sheet of the N-terminal domain (**Fig. 4a**). The latter loop contains the DFGWG motif (**Fig. 4b**).

**Figure 4.**
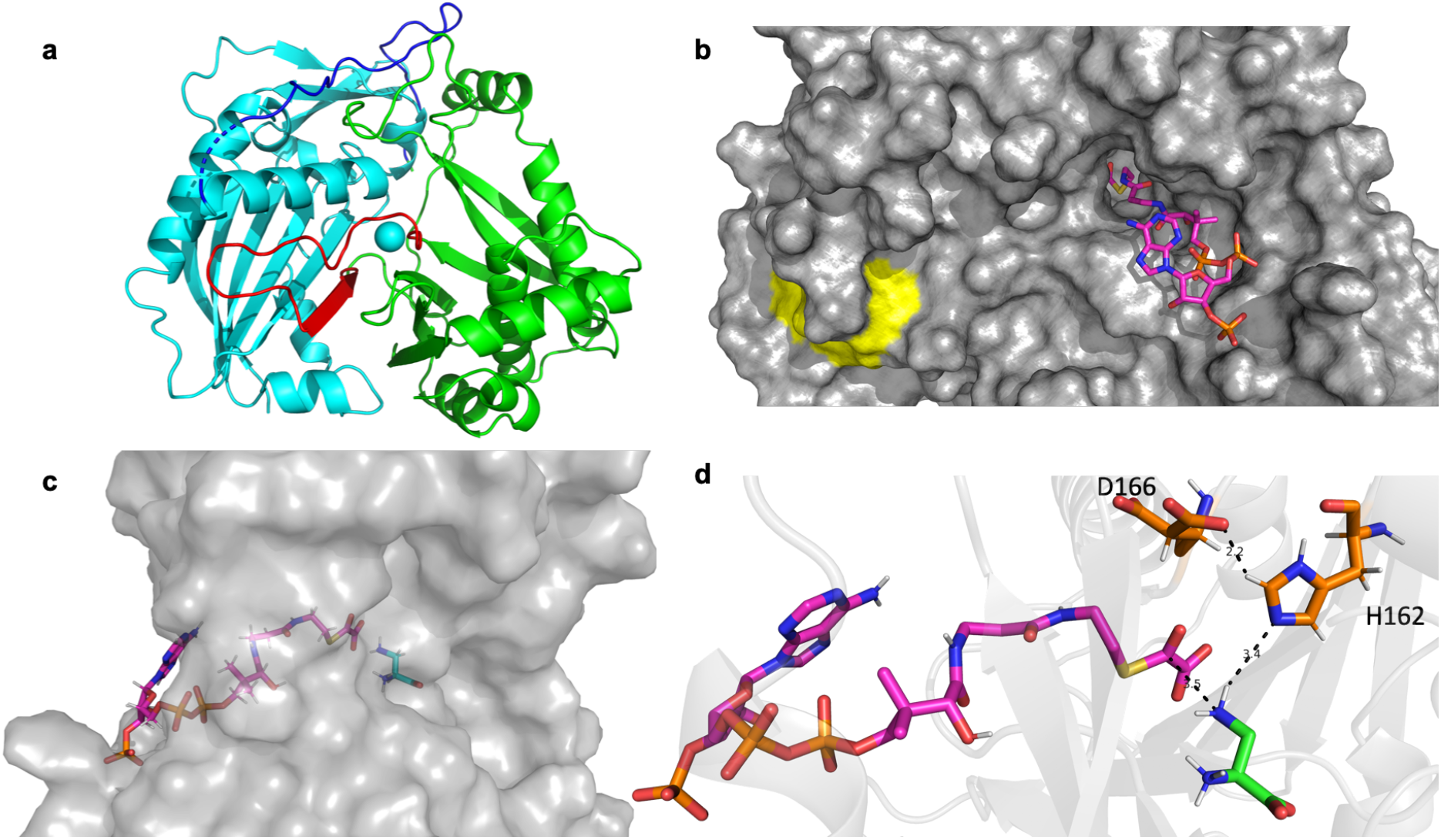
Crystal structure of BOS and docked substrates. **a.** Ribbon diagram of the crystal structure of BOS (PDB 6ZBS). The N-(cyan) and C-(green) terminal domains are connected by a loop (residues 183-209; blue) and a segment comprised of a β-strand and a loop (residues 371-393, red). **b**. Surface representation of BOS with bound oxalyl-CoA (magenta) docked into the active site. The conserved DFGWG motif is colored yellow **c.** Side view of BOS showing the orientation of oxalyl-CoA (magenta) and L-DAPA (cyan) docked into the active site. **d.** Close view of the active site showing the conserved catalytic residues His162 and Asp166 (orange) with H-bonds (dashed black lines), along with oxalyl-CoA (magenta) and L-DAPA (cyan) docked into the active site.

A search for structural homologs of BOS yielded six enzymes (**Fig. 2, S5**), all of which are plant hydroxycinnamoyl transferases (HCTs; **Fig. S7**). The main differences between BOS and its homologs are in loop lengths and conformations, and in the orientation of the openings of the active site tunnel (**Fig. S8 a,b**). The substrate binding site of HCTs is located in the interface between the two domains, inside a solvated tunnel that runs across the protein. Typically, the two tunnel openings are aligned with the tunnel (**Fig. S8 c,d**). By contrast, in BOS, the openings of the tunnel are angled to each other (**Fig. S8 e,f**).

To predict the mode of substrate binding by BOS and to gain insight into its catalytic mechanism, we docked its two substrates into the active site, using the structure of *Panicum virgatum* HCT ((*31*); PDB ID: 5FAL) as guide. We found that similar to other HCTs, the CoA substrate resides within the active site tunnel of BOS with its nucleotide moiety located at the entrance to the tunnel and its thiol moiety buried deep inside the tunnel (**Fig. 4b-d**). As expected, the other substrate, L-DAPA is positioned near the thiol moiety of oxalyl-CoA with which it interacts (**Fig. 4d**). Our docking studies showed that residue His162, of the conserved catalytic HXXXD motif, is located 3.5Å away from the β amine group of L-DAPA (**Fig. 4d**). Therefore, we suggest that similar to its role in other BAHD acetyltransferases (*26*), it acts to deprotonate the terminal-amino nitrogen of the substrate L-DAPA, enabling a nucleophilic attack on the carbonyl carbon of oxalyl-CoA. This, in turn, results in the formation of a tetrahedral intermediate that subsequently resolves to release CoA and the product, β-ODAP (**Fig. 4d, S9**). The role of the conserved Asp residue of this motif, Asp166, may be structural, as suggested (*26*), or may serve to activate His162 through hydrogen bonding (**Fig. 4d**). Our analysis also suggests that other residues, including Lys40, Asp284, Ser371, Ile369 and Lys401, may participate in substrate binding in the active site (**Fig. S10**).

During the process of this work we became aware of a study, published as a Ph.D. thesis (*32*), that pointed to the same gene in grass pea as a possible candidate for BOS. We compared the sequence of BOS that we identified, with that gene candidate, and found them to be identical.

Grass pea is a nutritious and robust crop. It is tolerant to both drought and flooding (2), and possesses a hardy and penetrating root system that enables it to grow on a wide range of soil types, including very poor soils and heavy clays (33). Therefore, it has the potential to improve global food security, especially under extreme climate conditions that are becoming more prevalent due to climate change (34). These attributes, however, are offset by the hazards associated with the production of the neurotoxin β-ODAP. The identification of BOS, along with its encoding gene, paves the way towards the generation of β-ODAP-free grass pea cultivars that would, hopefully, preserve the beneficial traits of the crop while being safe for consumption; adding a valuable, nutritious and robust resource to the world’s battery of food crops.

