## Supplementary Materials for "Bringing BOS to light: Uncovering the key enzyme in the biosynthesis of the neurotoxin β-ODAP in Grass Pea (*Lathyrus sativus* L.)"

### Supplementary Information[1]

#### Materials and Methods

##### Materials

Grass Pea (*Lathyrus sativus* L.*)* seeds were obtained from Fratelli Ingengnoli Milano© (https://www.ingegnoli.it) and from Plant World Seeds© (www.plant-world-seeds.com). Protease inhibitor cocktail for plant cells and for bacterial cells, Miracloth, HEPES, EDTA, ammonium sulfate, sodium phosphate monobasic and dibasic, sodium oxalate, magnesium chloride, coenzyme A sodium salt, 5,5'-dithio-bis-(2-nitrobenzoic acid) (DTNB), acetyl-CoA, malonyl-CoA, glutaryl-CoA, β-ODAP (SML1836), sodium tetraborate,$^{13}C_6$-Arginine hydrochloride and 2,3-diaminopropionic acid, benzonase, lysozyme, Amicon® Ultra-2 and 15 centrifugal filters were obtained from Sigma-Aldrich© and MERCK©. 6-aminoquinolyl-N-hydroxy-succinimyl carbamate (AQC) was synthesized following Cohen and Michaud (*1*). α-ODAP was obtained by partial thermal isomerization of β-ODAP(*2*). L-DAPA was obtained from AEchem Scientific©. Gelrite and Murashige & Skoog (MS) medium, including vitamins, were obtained from Duchefa©. RNeasy plant mini kit and DNeasy plant mini kit were obtained from Qiagen©. RNase-free DNase I was purchased from New England BioLabs©. Snakeskin dialysis tubing was obtained from ThermoFisher Scientific©. HiTrap Q FF 10/16, HiLoad 26/600 Superdex 200pg, HiTrap Blue HP and Superdex™ 200 increase 10/300 GL columns were obtained from GE Healthcare©. Zenix SEC-300 column was purchased from Sepax Technologies©. Synthetic DNA constructs were synthesized by Twist Bioscience©. 96-deepwell plates were obtained from Axygen®. 96-well ELISA plates were obtained from Greiner®. UPLC/MS grade solvents were used for all chromatographic steps.

##### Plant seed germination

Grass pea (*Lathyrus sativus*) seeds were surface sterilized by a brief wash with 70% ethanol and soaking in 3% sodium hypochlorite for 20min at constant, gentle (50 rpm) shaking. Then, they were rinsed four times with autoclaved DDW and kept in DDW O/N at 4ºC to synchronize their germination. For RNA extraction, the sterilized seeds were inoculated in a container supplemented with half-strength MS salts and 30 g/l sucrose and gelled with 0.3% gelrite at pH 5.8. For protein extraction the seeds were inoculated in plastic pots containing 250 $cm^3$ commercial potting mixture. Seeds were germinated under 12 h light (150 µmol $m^{-2}$ $s^{-1}$)/dark cycle at 25°C for one week.

##### Purification of β-ODAP synthetase from grass pea

Grass pea seeds (30gr) were soaked in sterile water for 3-4 days prior to processing. Sprouts (25gr) were harvested and washed with DDW prior to processing. Both seeds and sprouts were frozen in liquid nitrogen. Thawed seeds were ground using an electric hand blender. Sprouts were ground using a ceramic mortar and pestle. Ground tissue was resuspended in plant extraction buffer (50mM HEPES, pH 7.5; 1mM EDTA,

[1] Goldsmith et. al., "Bringing BOS to light: uncovering the key enzyme in the biosynthesis of the neurotoxin β-ODAP in Grass Pea"

pH 8.3; 250mM sucrose; protease inhibitor cocktail 1:250), homogenized using a glass homogenizer and pelleted (20,000xg, 1h, 4°C). The (viscous) supernatant was filtered through Miracloth and pelleted by ultracentrifugation (190,000xg, 1h, 4°C). It was then precipitated by ammonium-sulfate at 50% saturation (4°C, O/N) and re-pelleted (20,000xg, 1h, 4°C). Pellets were resuspended in buffer A (50mM HEPES, pH 7.5; 1mM EDTA, pH 8.3; 50mM NaCl), dialyzed extensively (10K MWCO, 22 mm) against buffer A and the solution was then pelleted (20,000xg, 1h, 4°C). The supernatant was loaded onto a HiTrap-Q-FF-10/16™ column connected to an AKTA pure 25M HPLC (GE Healthcare©) and eluted using a gradient from buffer A to buffer B (50mM HEPES, pH 7.5; 1mM EDTA, pH 8.3; 1M NaCl). Fractions exhibiting BOS activity were pooled, concentrated and washed with buffer A using Amicon® Ultra-15 centrifugal filters (10kDa MWCO). The concentrates were loaded onto a HiLoad 26/600 Superdex 200pg column and eluted using buffer C (50mM HEPES, pH 7.5; 1mM EDTA, pH 8.3; 150mM NaCl). Eluted fractions exhibiting BOS activity were again pooled, concentrated and washed with buffer D (20mM Na-phosphate, pH 7.5; 1mM EDTA, pH 8.3), and loaded onto a HiTrap Blue HP (5ml) column. BOS-containing fractions were eluted using a gradient from buffer D to buffer E (20mM Na-phosphate, pH 7.5; 1mM EDTA, pH 8.3; 2M NaCl). Fractions exhibiting BOS activity were pooled, washed with buffer A and re-concentrated. The concentrates were loaded on a Superdex™ 200 increase 10/300 GL and eluted using buffer C or loaded onto a Zenix sec 300 and eluted with PBS buffer. Fractions exhibiting BOS activity were then pooled, rinsed with 50mM NaCl, concentrated using Amicon® Ultra 2mL centrifugal filters (10kDa MWCO) and sent for peptide sequencing.

**Synthesis of oxalyl-CoA using oxalyl CoA from grass pea**

The gene encoding oxalyl CoA-synthetase (OCS) was amplified from extracted genomic DNA of *L. sativus*, cloned into a bacterial expression vector, expressed in *E.coli* cells and purified (see ref (*3*)). Purified OCS (0.25μM) was added to a reaction solution (125mM HEPES, pH 7.5; 2mM $MgCl_2$; 10mM ATP) containing oxalic acid (5mM) and coenzyme A (1.5mM) and incubated at 37°C for 1h. The reaction mix was sampled before the addition of OCS and at the end of the incubation. The samples, diluted two-fold, were mixed with an equal volume of buffered DTNB solution (50mM HEPES, pH 7.5; 1mM DTNB) and incubated for 15min at R.T. The amount of unreacted coenzyme A was derived by comparing the absorbance of the sample solutions at 412nm, before and after incubation with OCS. Typically, under these conditions, more than 95% of coenzyme A reacted to form oxalyl-CoA.

**L-DAPA elimination assay**

The activity of β-ODAP synthetase was determined in purified protein fractions as previously described (*4*). Oxalyl-CoA was freshly prepared *in-vitro* as described above. Samples of the oxalyl-CoA reaction mixture (oxalyl-CoA 1.5mM, 40μl) were mixed with buffered DAPA solution (50mM HEPES, pH 7.5; 0.7mM, 30μl DAPA) and purified grass pea protein fractions (30μl). The fractions were incubated at 37°C for 15-30 min in 96-deepwell plates and then mixed with an OPT reagent solution (50mM boric acid, pH 9.9; 60mM β-mercaptoethanol; 7.5mM, 400μl o-phthalaldehyde). DAPA concentrations were assayed by measuring solution absorbance at 422nm in duplicates (200μl) in 96-well ELISA plates.

**Grass pea cDNA library preparation an sequencing**
Total RNA from five-week old grass pea plants was isolated using RNeasy plant mini kit and residual DNA was removed with RNase-free DNase I. RNA was quantified using a Qubit 3 fluorometer and an RNA broad-range kit (Invitrogen©). RNA integrity was assessed using an Agilent 2100 Bioanalyzer. Only RNA with an integrity number (RIN) greater than 8 was used for SMRTbell library preparation. An Iso-Seq library was prepared according to the Isoform sequencing protocol, using the Clontech SMARTer PCR cDNA Synthesis kit as described by Pacific Biosciences with the following modifications: One microgram of RNA, combined from two different tissues, was used as input in the Clontech SMARTer reaction. PCR cycle optimization was performed by amplification of first-strand cDNA by 10 to 18 cycles using PrimeSTAR GXL polymerase (Takara). Large-scale PCR was performed by 11 amplification cycles to generate double-stranded cDNA for SMRTbell library construction. The amplified cDNA was used to build SMRTbell template libraries according to the Iso-Seq protocol. One SMRT cell was sequenced using the PacBio Sequel platform.

**Iso-Seq data analysis**
Raw data were processed using SMRTlink 4.0 software. Circular consistency sequences (CCSs) were generated into full-length reads, which were clustered into full-length non-chimeric transcripts by ISOseq3. From these transcripts, high quality (HQ) isoforms ($Q_{score}$>30) were selected for further analysis. Since there was no reference genome for grass pea, the Coding Genome Reconstruction Tool (Cogent) was used to generate a fake coding genome on which Cupcake ToFU was employed to remove redundant sequences. This pipeline yielded a non-redundant, high likelihood gene assembly. The 'Dumb' algorithm was used to find the longest open reading frames (ORFs). We then used the 'ANGEL'(*5*) algorithm to predict the most likely ORFs based on length and coding potential of each given sequence. Gene ontology (GO) annotation analysis was performed using Blast2GO (http://www.blast2go.org/) version 2.5.0. After gene-ID mapping, GO term assignment, annotation augmentation and generic GO-slim process, the final annotation file was produced. The data were submitted as BioProject ID PRJNA662941 (http://www.ncbi.nlm.nih.gov/bioproject/662941).

**Proteomic analysis**
Protein fractions were subjected to in-solution tryptic digestion, using the suspension trapping (S-trap) as described (*6*). Briefly, protein concentration was measured using the BCA assay (Thermo Scientific, USA). Protein fractions were supplemented with SDS (5% final concentration) in 50mM Tris-HCl, reduced with 5 mM dithiothreitol and alkylated with 10mM iodoacetamide in the dark. Samples were loaded onto S-Trap microcolumns (Protifi, USA) according to the manufacturer's instructions, and the columns were washed with 90:10% methanol/50mM ammonium bicarbonate. Samples were then digested with trypsin (1:50 trypsin/protein) for 1.5 h at 47°C. The digested peptides were eluted using 50mM ammonium bicarbonate. Trypsin was added to this fraction that was further incubated O/N at 37°C. Two more elutions were made using 0.2% formic acid and 0.2% formic acid in 50% acetonitrile. The three elutions were pooled together and vacuum-centrifuged to dryness. Samples were kept at −80°C until further analysis. Dry digested samples were dissolved in 97:3% $H_2O$/acetonitrile + 0.1% formic acid. Peptides were separated by chromatography using nano-Ultra Performance Liquid Chromatography (10 kpsi nanoAcquity; Waters, Milford, MA). The mobile phase was: A) $H_2O$ + 0.1% formic acid and B) acetonitrile +

0.1% formic acid. Desalting of the samples was performed online using a reversed-phase Symmetry C18 trapping column (180 µm internal diameter, 20 mm length, 5 µm particle size; Waters). Peptides were separated using a T3 HSS nano-column (75 µm internal diameter, 250mm length, 1.8µm particle size; Waters) at 0.35µL/min and were eluted into the mass spectrometer using the following gradient: 4% to 30%B in 50 min, 30% to 90%B in 5 min, maintained at 90% for 5min and then back to initial conditions. The nanoUPLC was coupled online through a nanoESI emitter (10 µm tip; New Objective; Woburn, MA, USA) to a quadrupole Orbitrap mass spectrometer (Sample BOS1: Q Excactive Plus, Samples BOS2-3: Fusion Lumos, BOS4: Q Exactive HFX, Thermo Scientific) using a FlexIon nanospray apparatus (Proxeon). Data were acquired in data-dependent acquisition (DDA) mode, using the Top10 method. MS1 resolution was set to 120,000 (at 200m/z), mass range of 375-1650m/z, AGC of 3e6, and the maximum injection time was set to 60msec. MS2 resolution was set to 15,000, quadrupole isolation 1.7m/z, AGC of 1e5, dynamic exclusion of 20sec and maximum injection time of 60 msec. Raw data were processed with Proteome Discoverer V2.2, and searched using SequestHT (*7*) and Mascot (*8*) engines against the translated transcriptome as the protein database, appended with common lab protein contaminants. Enzyme specificity was set to trypsin and up to two missed cleavages were allowed. Fixed modification was set to carbamidomethylation of cysteines and variable modifications were set to oxidation of methionines, protein N-terminal acetylation and deamidation of glutamines and asparagines. Peptide precursor ions were searched using a maximum mass deviation of 10ppm and fragment ions using maximum mass deviation of 0.02Da. Peptide and protein identifications were filtered at an FDR of 0.01 using the decoy database strategy.

**β-ODAP synthetase activity assays in *E.coli* cell lysates**

Selected grass pea protein sequences, codon-optimized for *E.coli* expression, were synthesized and cloned into a pET-His-SUMO expression vector. The vector was constructed by transferring the His14-bdSUMO cassette from the expression vector (designated K151), generously obtained from Prof. Dirk Görlich from the Max-Planck-Institute, Göttingen (*9*), into the expression vector pET28-TevH (*10*). Expression plasmids were transformed into electrocompetent BL21/DE3 cells harboring a chaperone-expressing plasmid pGJKE8 (Takara©) and plated on LB agar plates with 35 µg/ml chloramphenicol, 50 µg/ml kanamycin and 1% w/v glucose. Following O/N incubation at 37°C, 3 individual colonies from each gene transformation were randomly picked into 96-Deep-Well plates (Axygen®) containing growth media (2YT + Cap 35µg/ml + Kana 50µg/ml + Arabinose 0.5mg/ml; 500µl/well) and grown O/N (R.T, shaking 1000RPM). The resulting cultures were used to inoculate (1:100 dilution) 0.5 ml growth media in 96-deepwell plates and grown at R.T to $OD_{600}$nm ≈ 0.6. Cultures were induced by IPTG (1mM), and grown O/N (R.T, shaking 1000RPM). Following growth, cells were pelleted (4000RPM, 4°C, 20min), frozen at -80°C, and thawed by the addition of cell lysis buffer (50mM HEPES, pH 7.5; 100mM NaCl; 2mM $MgCl_2$; 0.6mg/ml lysozyme; 10U/ml benzonase nuclease®; bacterial protease inhibitor cocktail without EDTA 1:50). Cells were lysed by shaking (1h, 37°C, 1200 RPM), and the lysate was clarified by centrifugation (4000RPM, 4°C, 30min). Clarified cell lysate (20µl), oxalyl-CoA reaction mix (1.5mM; 20µl) and a buffered DAPA solution (50mM HEPES, pH 7.5; 0.7mM, 20µl, DAPA) were mixed in 96-well ELISA plates and incubated (15min, 37°C). Then, 200µl of OPT reagent solution (50mM boric acid, pH 9.9; 60mM β-mercaptoethanol; 7.5mM o-phthalaldehyde) were added into each well, incubated for 10min, at R.T and the solution absorbance was read at 422nm using a plate reader (BioTeck©). Negative control samples contained cell lysis buffer only and positive control samples contained grass pea lysate (20μl).

**Cloning, expression and purification of BOS**
Expression of BOS was performed using the expression vector pET28-bdSumo described above. A 5 L culture of BL21(DE3) was induced with 200μM IPTG and grown O/N at 15°C. The cells were harvested and lysed by cell disrupter (Constant Systems©) in lysis buffer (50mM Tris, pH 7.5, 0.1M NaCl, 1mM DTT, 2mM $MgCl_2$, 8% glycerol) containing 200KU/100 ml lysozyme, 20μg/ml DNase, 1mM phenylmethylsulfonyl fluoride (PMSF) and protease inhibitor cocktail. Following centrifugation, 1% Triton was added to the sup and the lysate was incubated with 5ml rinsed Ni beads (Adar Biotech©) for 1h at 4°C. The beads were washed 4 times with 50ml lysis buffer. The enzyme was eluted from the beads by incubation with 5ml lysis buffer containing 0.4mg bdSumo protease for 2h at R.T. The solution containing cleaved BOS was collected and additional 5ml cleavage buffer was added for 2h at R.T. The two elutions were combined, concentrated and applied to a size exclusion (SEC) column (HiLoad 16/60 Superdex75 prep-grade, GE Healthcare©) equilibrated with 50mM Tris pH 7.5, 0.1M NaCl, 2mM $MgCl_2$, 8% glycerol. Pure BOS, migrating as a single peak at 58ml, was pooled, concentrated to 28 mg/ml and frozen in aliquots at -80°C.

**Determination of catalytic efficiency parameters**
*In-situ* synthesized oxalyl-CoA (6mM), acetyl-CoA (2mM), malonyl-CoA (2mM) or glutaryl-CoA (2mM) were mixed with purified BOS (0.1μM) and varying concentrations of L-DAPA (0.09-4.8mM) in buffer A (50mM HEPES, pH 7.5; 1mM EDTA, pH 8.3; 50mM NaCl) at room temperature. Reaction mixtures were sampled at different time intervals during the first 5min of the reaction and the reactions were quenched by mixing into an OPT solution (50mM boric acid, pH 9.9; 60mM β-mercaptoethanol; 7.5mM o-phthalaldehyde) at a 1:9 (v/v) ratio. Solution absorbance was read at 422nm in 96-well plates using an ELISA plate reader (BioTek©). Results obtained with oxalyl-CoA were fit to a Michaelis-Menten equation to derive $k_{cat}$ and $K_M$ of BOS. Results obtained with other CoA substrates were fit to the linear part of the Michaelis-Menten curve to derive the value of $k_{cat}/k_M$.

**LC-MS analysis of β-ODAP and L-DAPA concentrations**
Sample derivatization was performed as described (*1, 11*). Briefly, a 10μL sample aliquot or amino acid standard solution were mixed with $^{13}C_6$-Arg (1μL, 60μM) as internal standard with 69μL of 0.15M sodium borate solution, pH 8.8. Solutions were derivatized using 20μL of 6-aminoquinolyl-N-hydroxy-succinimyl carbamate in acetonitrile (2.7mg/mL) by incubation at 55°C for 10 min. The reaction mixtures were cooled and placed in nanofilter vials (Thomson, 0.2μm PES) for LC-MS/MS analysis, using an Acquity I-class UPLC system (Waters©) and a Xevo TQ-S triple quadrupole mass spectrometer (Waters©) equipped with an electrospray ion source. LC was performed using a 100 × 2.1-mm i.d., 1.8μm UPLC HSS T3 column (Waters©) with mobile phases A (0.1% aqueous formic acid) and B (0.1% formic acid in acetonitrile) at a flow rate of 0.4 ml/min and column temperature of 35°C. The gradient was as follows: 2%B for 1min, then a linear increase to 30% B (7min), then an increase to 100%B (0.5 min), 100%B for 1.5 min, then reduce to 2% B (0.5 min), and 2% B for

1.5 min. Samples were kept at 4°C and automatically injected. Sample volume was 1μl. Mass spectrometry was performed in positive ion mode, with capillary voltage of 3kV, cone voltage of 23V, desolvation temperature of 350°C, source temperature of 120°C, desolvation gas flow of 500L/h, and collision gas flow of 0.10ml/min. The following MS/MS transitions were monitored: m/z 347.0 → 171.0 for α- and β-ODAP (retention time 4.02 and 3.81min, respectively), 275.1 → 171.0 and 223.1 → 171.0 for DAPA (as 1xAQC and 2xAQC adducts, respectively) and 351.0 → 171.0 for $^{13}C_6$-Arg as internal standard.

**Crystallization, data collection and refinement**

Purified BOS was crystallized using the hanging drop vapor diffusion method and a Mosquito robot (TTP LabTech©) at 19°C, utilizing the precipitants 0.1M $MgCl_2$ and 10% polyethylene glycol (PEG) 6,000 in 50mM HEPES (pH 7.0). BOS crystals formed in the tetragonal space group $P4_32_12$, with one monomer per asymmetric unit and diffracted to 2.35Å resolution. Data collection was performed under cryo conditions (100K), in-house, using a Rigaku RU-H3R X-ray instrument. All diffraction images were indexed and integrated using the Mosflm program (*12*), and the integrated reflections were scaled using the SCALA program (*13*). Structure factor amplitudes were calculated using TRUNCATE (*14*) from the CCP4 program suite. The structure of BOS was solved by molecular replacement with the program PHASER (*15*), using the distantly related (23% sequence identity) hydroxycinnamoyltransferase (HCT) structure from *Sorghum bicolor* (PDB-ID 4KE4). All steps of the atomic refinements were performed with the PHENIX.refine, Parallel PHENIX.phaser programs (*16*). The model was built into 2mFobs- DFcalc, and mFobs - DFcalc maps using COOT (*17*), optimized using PDB_REDO (*18*), and evaluated using MOLPROBIDITY (*19*). The crystal structure was deposited in the PDB with the accession number: 6ZBS.

**Definition of BAHD family members and phylogenetic analysis**

Database searches (BlastP) were performed at the NCBI website against the non-redundant (NR) database with default parameters. The full complement of BAHD and *Arabidopsis* protein sequences were compared using TBlastN in a local installation (version 2.6.0+ (*20*)) against the PacBio transcripts. Hits with an E-score $\leq 10^{-30}$ were taken for further analysis. DNA sequence redundancy was examined using the Sequencher software (version 5.4.6, GeneCodes Corporation©, Ann Arbor, MI, USA). Domain analysis was performed at NCBI, using the Batch-CDD Search tool (*21*). Multiple alignment was performed with ClustalW (version 2.1 (*22*)). The full complement of sequences from *Populus trichocarpa* was added (*23*). Ninety-one sequences with known function were taken from refs: (*24, 25*) and 67 additional, non-redundant sequences from other species were added. Alignments were performed with ClustalW. Phylogenetic analysis was performed with Neighbor-Joining in ClustalW, Phylip (version 3.697 (*26*)) and PhyML online (version 3 (*27*)), using default parameters. The PhyML tree was visualized with iTOL (*28*).

**Molecular docking studies**

The three-dimensional crystal structure of BOS (solved here; PDB ID: 6ZBS) was optimized prior to docking using the Protein Preparation Wizard in Schrödinger Maestro Suite 2019 (Schrödinger Suite 2019-2 Protein Preparation Wizard; Epik, Schrödinger, LLC, New York, NY, 2019; Impact, Schrödinger, LLC, New York, NY, 2018; Prime, Schrödinger, LLC, New York, NY, 2018). Inconsistencies in the structure, such as missing side-chains and hydrogens, incorrect bond orders or the orientation of the different functional groups of the amino acids were rectified during the optimization process (*29*) and the prepared protein was used for GLIDE Docking (*30-32*). The structures of the oxalyl-CoA, L-DAPA and β-ODAP ligands were prepared prior to docking using the LigPrep application in Schrödinger Maestro Suite 2019 (LigPrep, Schrödinger, 2019-2). LigPrep was used to perform the conversion of structures from 2-dimensional to 3-dimensional, correction of improper bond distances, bond orders, generation of ionization states and energy minimization processes. The ligand structures were then used for rigid protein-ligand docking using the GLIDE docking application (Glide, Schrödinger, 2019-2). Default settings were selected for the docking protocol with Standard Precision (SP) scoring function mode. For the oxalyl-CoA docking procedure, the crystal structure of PvHCT in complex with CoA and p-coumaroyl-shikimate (PDB code 5FAL) was superimposed on the BOS structure and the CoA ligand was used to generate the enzyme grid which was then used for the docking protocol. For the L-DAPA and β-ODAP ligands, the best oxalyl-CoA docking result was selected and truncated to a four carbon chain that was used as a dummy ligand for the grid-generation. The poses were ordered based on the GLIDE docking score.

**Supplementary Table S1: Summary of PacBio long-read cDNA sequencing**

| | |
|---|---|
| Number of reads of insert | 15,046,014 |
| Number of read bases of insert | 24,422,044,159 |
| Average length of ROI[a] (bp) | 1,623 |
| Number of non-full-length PacBio reads | 65,228 |
| Number of FLNC[b] PacBio reads | 473,833 |
| Average length of FLNC PacBio reads (bp) | 1,764 |
| Number of polished high-quality reads | 39,350 |
| Number of polished low-quality reads | 183 |

[a.] ROI – reads of insert.
[b.] FLNC – Full-length, non-concatemer

**Supplementary Table S2. Number of proteins identified in samples***

| | **BOS-1** | **BOS-2** | **BOS-3** | **BOS-4** |
|---|---|---|---|---|
| Total number of proteins found | 1969 | 1476 | 743 | 247 |
| Proteins shared with some samples | 1197 | 1182 | 479 | 22 |
| Proteins shared between all samples | 222 | | | |
| Unique proteins | 550 | 72 | 42 | 3 |

*Proteins were identified using the SequestHT (*7*) and Mascot (*8*) search engines, searching against the translated transcriptome as the protein database.

#### Supplementary Table S3: Data collection and refinement statistics for BOS

| | |
|---|---|
| **Data Collection** | |
| PDB code | 6ZBS |
| Space group | $P4_32_12$ |
| Cell dimensions: | |
| a,b,c (Å) | 66.20 66.20 226.38 |
| α,β,γ (˚) | 90, 90, 90 |
| No. of copies in a.u. | 1 |
| Resolution (Å) | 45.84-2.35 |
| Upper resolution shell (Å) | 2.434-2.35 |
| Unique reflections | 21,933 (2,130) |
| Completeness (%) | 99.83 (99.77) |
| Multiplicity | 19.3 (21.1) |
| Average I/σ(I) | 49.73 (5.75) |
| R-pim | 0.165 (0.2717) |
| CC1/2 | 0.75 (0.824) |
| **Refinement** | |
| Resolution range (Å) | 45.84-2.35 |
| No. of reflections (I/σ(I) > 0) | 21,906 |
| No. of reflections in test set | 1,112 |
| R-working / R-free | 0.2059 / 0.2590 |
| No. of protein atoms | 3324 |
| No. of water molecules | 111 |
| Overall average B factor (Å$^2$) | 43.73 |
| Root mean square deviations: | |
| - bond length (Å) | 0.006 |
| - bond angle (°) | 0.85 |
| **Ramachandran Plot** | |
| Most favored (%) | 95.45 |
| Additionally allowed (%) | 3.83 |
| Disallowed (%) | 0.72 |

* Values in parentheses refer to the data of the corresponding upper resolution shell.

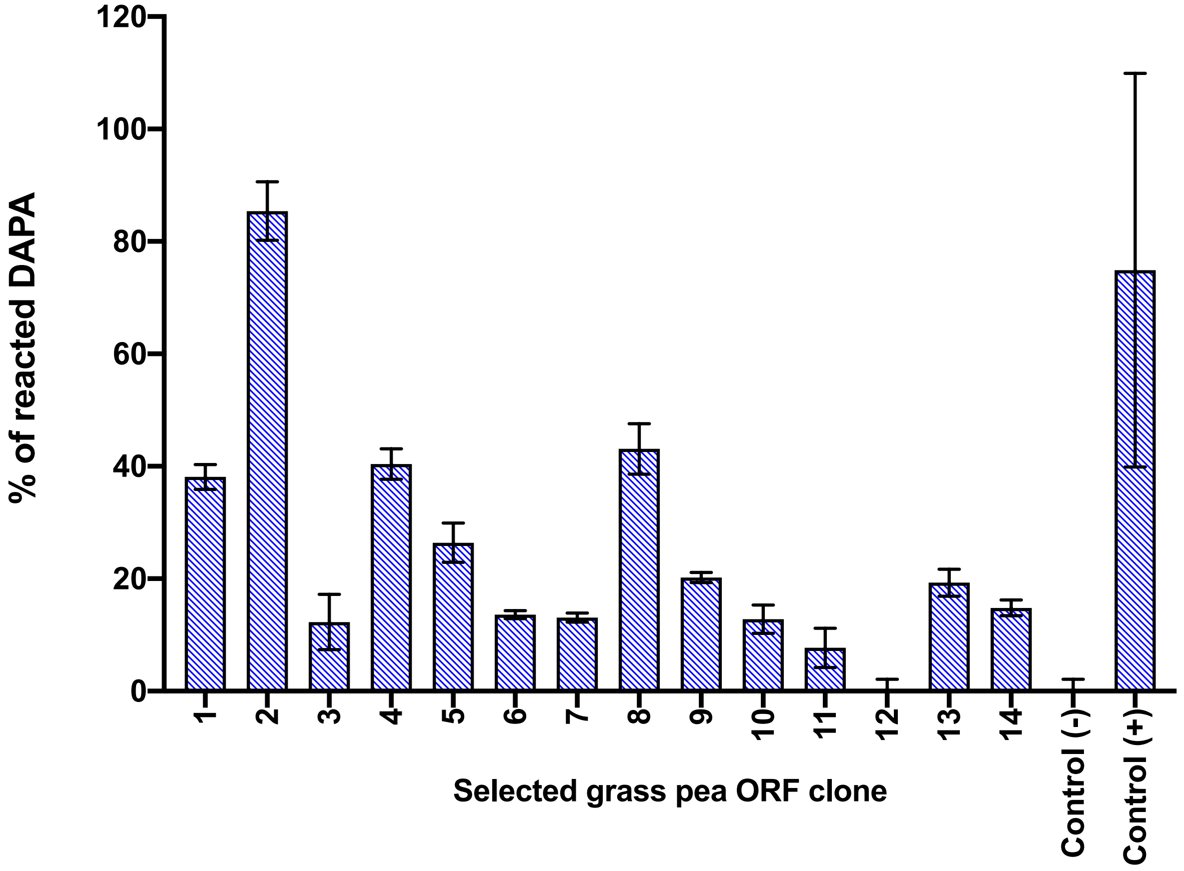


**Supplementary Figure S1. DAPA elimination activity of grass pea ORFs expressed in *E.coli* cells.** Selected ORFs from the grass pea transcriptome were cloned, transformed and expressed in *E.coli* cells. DAPA elimination was assayed in cell lysates as described in Materials and Methods. Negative controls contained lysis buffer only. Positive controls contained grass pea leaf lysates. Error bars denote the standard deviation of 3 replicate measurements.

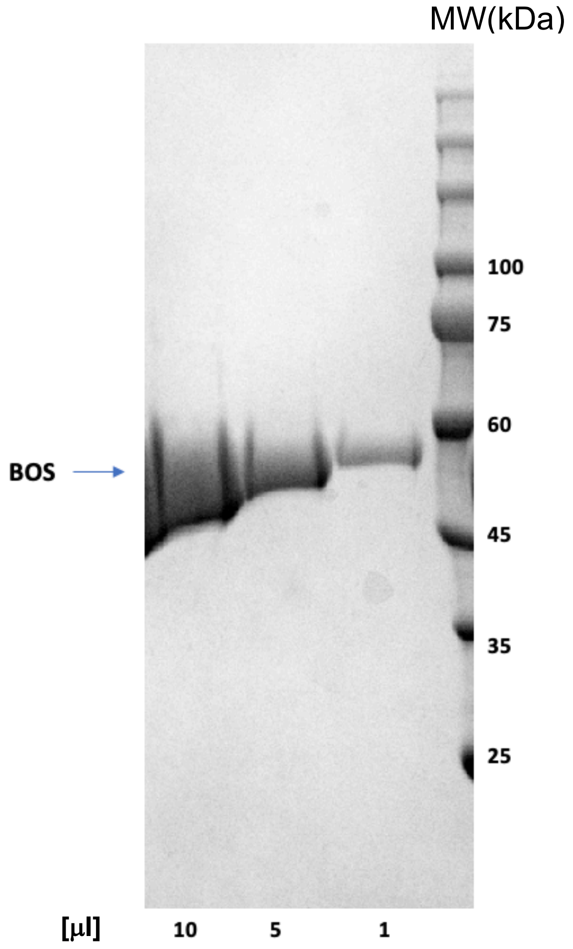


**Supplementary Figure S2. SDS-PAGE gel of purified BOS.**
Increasing amounts of purified BOS protein were run on a 12% SDS-PAGE gel and stained with Coomassie blue. Right lane – prestained molecular weight markers.

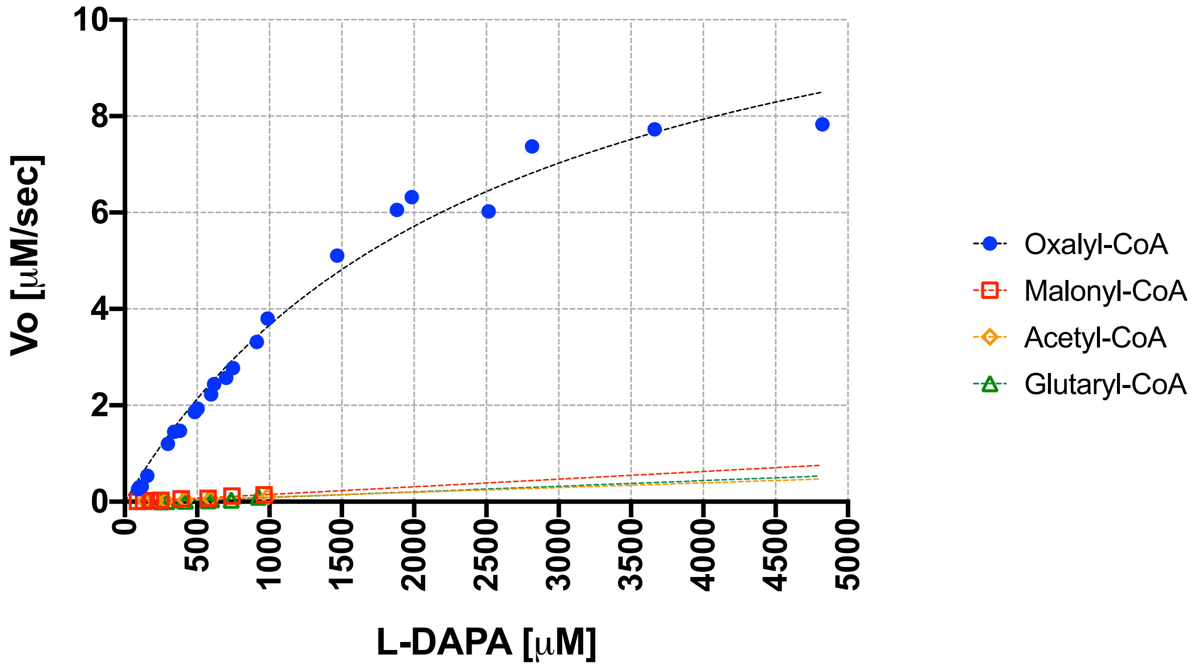


**Supplementary Figure S3. Kinetics of CoA substrate ligation to L-DAPA catalyzed by BOS.** Shown is a Michaelis-Menten plot for the β-ODAP synthetase activity of BOS. *In-situ* generated oxalyl-CoA (6mM) or commercially obtained acetyl-CoA (2mM), malonyl-CoA (2mM) or glutaryl-CoA (2mM) were mixed with different concentrations of L-DAPA (0.09-4.8mM) and purified BOS (0.1μM) at room temperature. Initial velocities ($V_0$) were obtained by measuring residual L-DAPA concentrations at different time points. The analysis of the data and the calculation of the kinetic constants by nonlinear regression analysis were performed using GraphPad Prism 8.4.3 for Mac OS X, GraphPad Software, San Diego, California USA, www.graphpad.com.

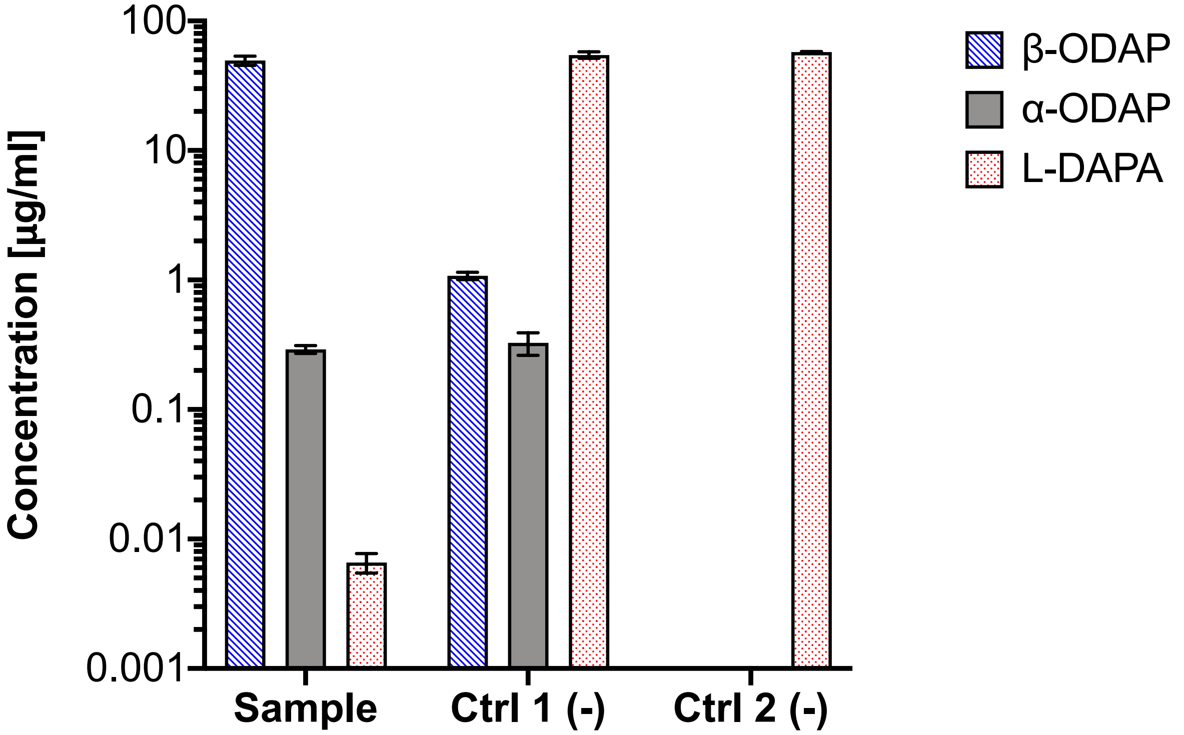


**Supplementary Figure S4. BOS catalyzes the formation of β-ODAP from oxalyl-CoA and L-DAPA.** Shown are the concentrations of β-ODAP, α-ODAP and L-DAPA determined using LC-MS from *in-vitro* reaction samples and controls. The complete reaction mixture (sample) contained a buffered solution of purified BOS (1μM), L-DAPA (480μM), and oxalyl-CoA (1.8mM). The Ctrl 1 (-) reaction mixture contained only L-DAPA (480μM) and oxalyl-CoA (1.8mM). The Ctrl 2 (-) reaction mixture contained only L-DAPA (480μM). Reaction mixtures were incubated (1h, 37ºC), sampled and examined by LC-MS (**Materials and Methods**). Error bars denote the standard deviation from 5 replicate samples**.**

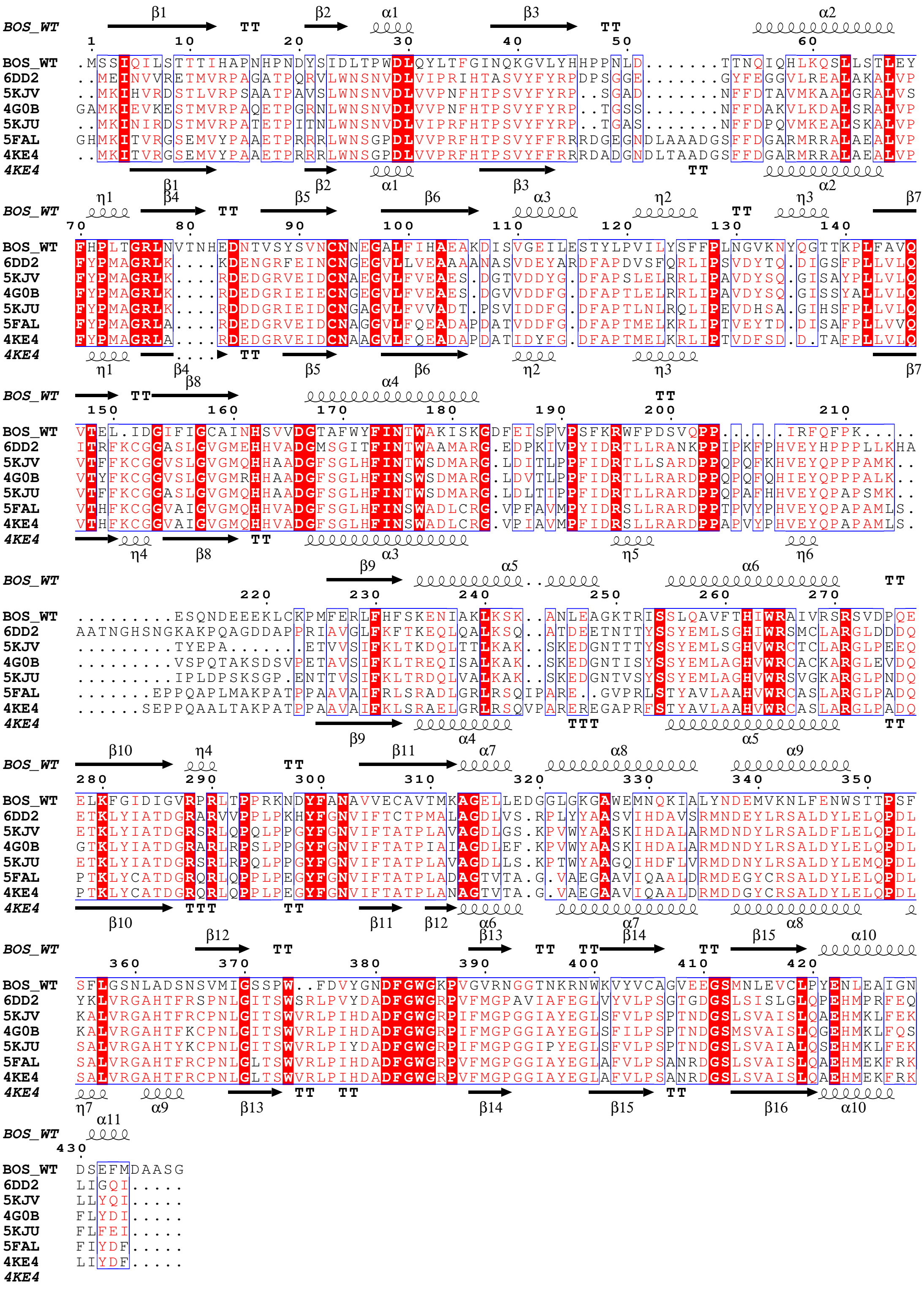

**Supplementary Figure S5. Secondary structure-based sequence alignment of BOS and HCT homologs.** The sequences of CoA:shikimate hydroxycinnamoyl transferases (HCTs) from different species were aligned to that of BOS. The structures are labeled according to their PDB codes: *Selaginella moellendorffii* (6DD2); *Plectranthus scutellarioides* (5KJV); *Coffea canephora* (4G0B*); Arabidopsis thaliana* (5KJU); *Panicum virgatum* (5FAL); *Sorghum bicolor* (4KE4). BOS secondary structure elements are labeled above the sequences and those of *Sorghum bicolor* (4KE4) - at the bottom. α-helices and η-helices are depicted as spirals, and β-strands are depicted as arrows. The residues conserved in all proteins are shown in red blocks. The multiple sequence alignment was produced using MultAlin (*33*) (Multiple sequence alignment with hierarchical clustering), and the figure was created using ESPript (*34*)

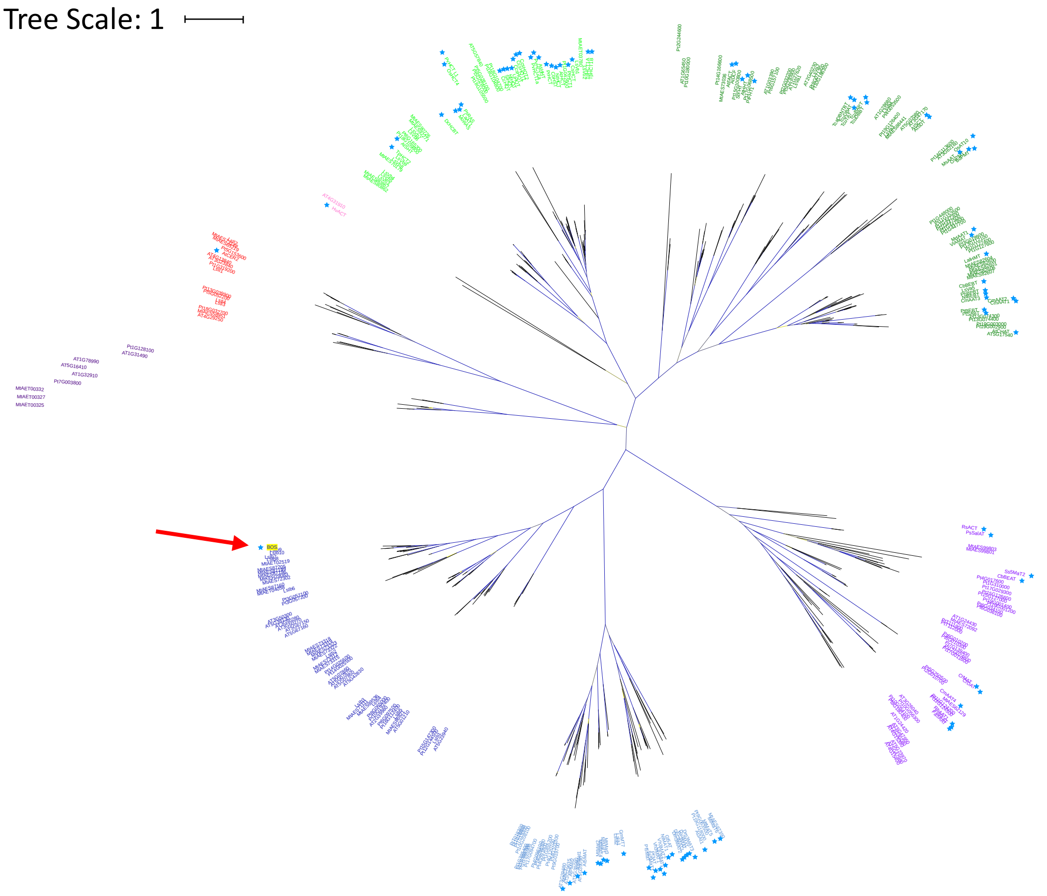


**Supplementary Figure S6. Phylogenetic tree with support coloring.** The phylogenetic tree from **Figure 3** is shown in unrooted form with support values colored in a range from blue (1) to yellow (0). Sequences names are colored according to clade: IIIb (dark purple), II (red), IV (pink), Vb (light green), Va (dark green), IIIa (light purple), Ia (light blue), Ib (dark blue). Sequences with known function are starred; BOS has a yellow background and is marked by a red arrow.

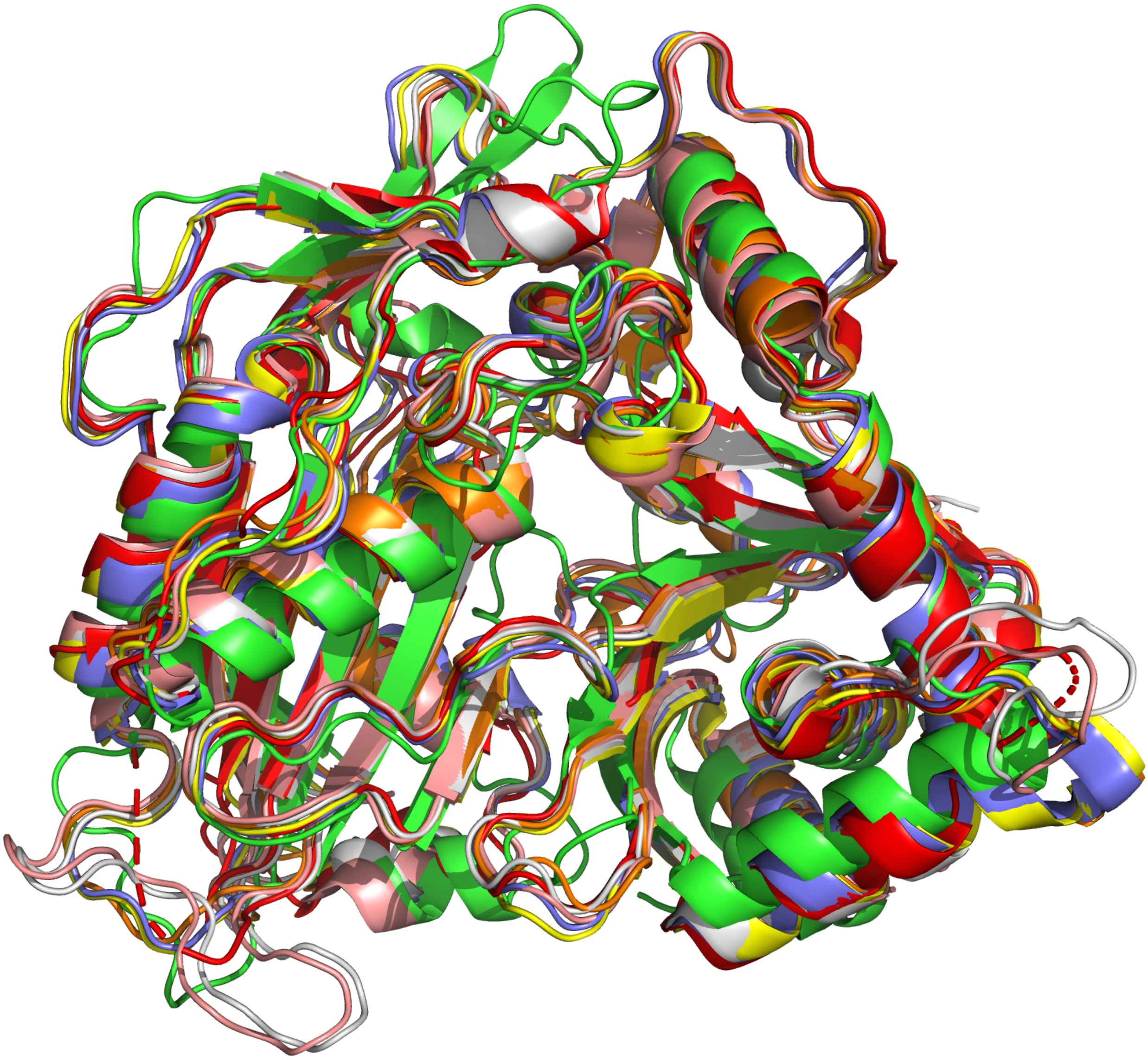

**Supplementary Figure S7. Structural overlay of BOS with five homologs.** The crystal structure of BOS (green) was aligned with the structures of six hydroxycinnamoyl transferases from: *Selaginella moellendorffii* (PDB ID: 6DD2, red), *Arabidopsis thaliana* (PDB ID: 5KJU, orange), *Coffea canephora* (PDB ID: 4G0B, yellow), *Panicum virgatum* (PDB ID: 5FAL, pink), *Sorghum bicolor* (PDB ID: 4KE4, gray), and *Plectranthus scutellarioides* (PDB ID: 5KJV, purple). Structures are presented as cartoon models without bound ligands. The figure was created using the PyMol molecular graphics system, version 2.4, Schrödinger, LLC.

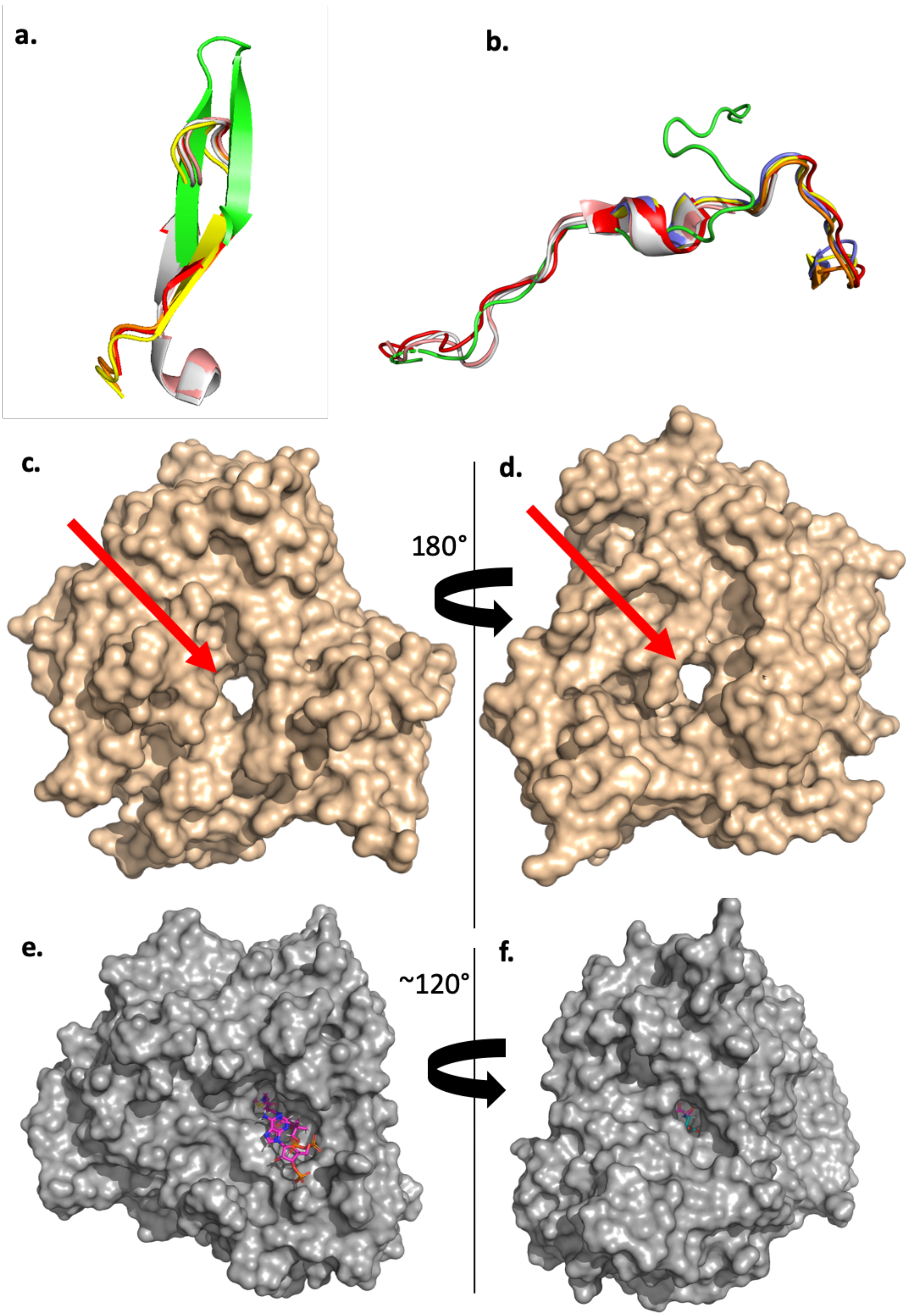
a.
b.
c.
d.
180°
e.
f.
~120°

**Supplementary Figure S8. Unique structural features of BOS.**
**a.** Loop residues 76-93 of BOS (green) are more extended than the other 6 HCT homologs. **b.** The conformation of the loop comprising residues 199-210 of BOS (green) is different from its conformation in other HCTs. **c,d.** Typically the openings of the active site tunnel in HCT homologs are aligned one opposite the other as in the HCT from *Sorghum bicolor* (PDB ID: 4KE4, gold). **e,f.** The openings of the active site tunnel in BOS (gray) are bent relative to each other. Oxalyl-CoA (magenta) and L-DAPA (cyan) were docked into the active site. The figure was created using the PyMol molecular graphics system, version 2.4, Schrödinger, LLC.

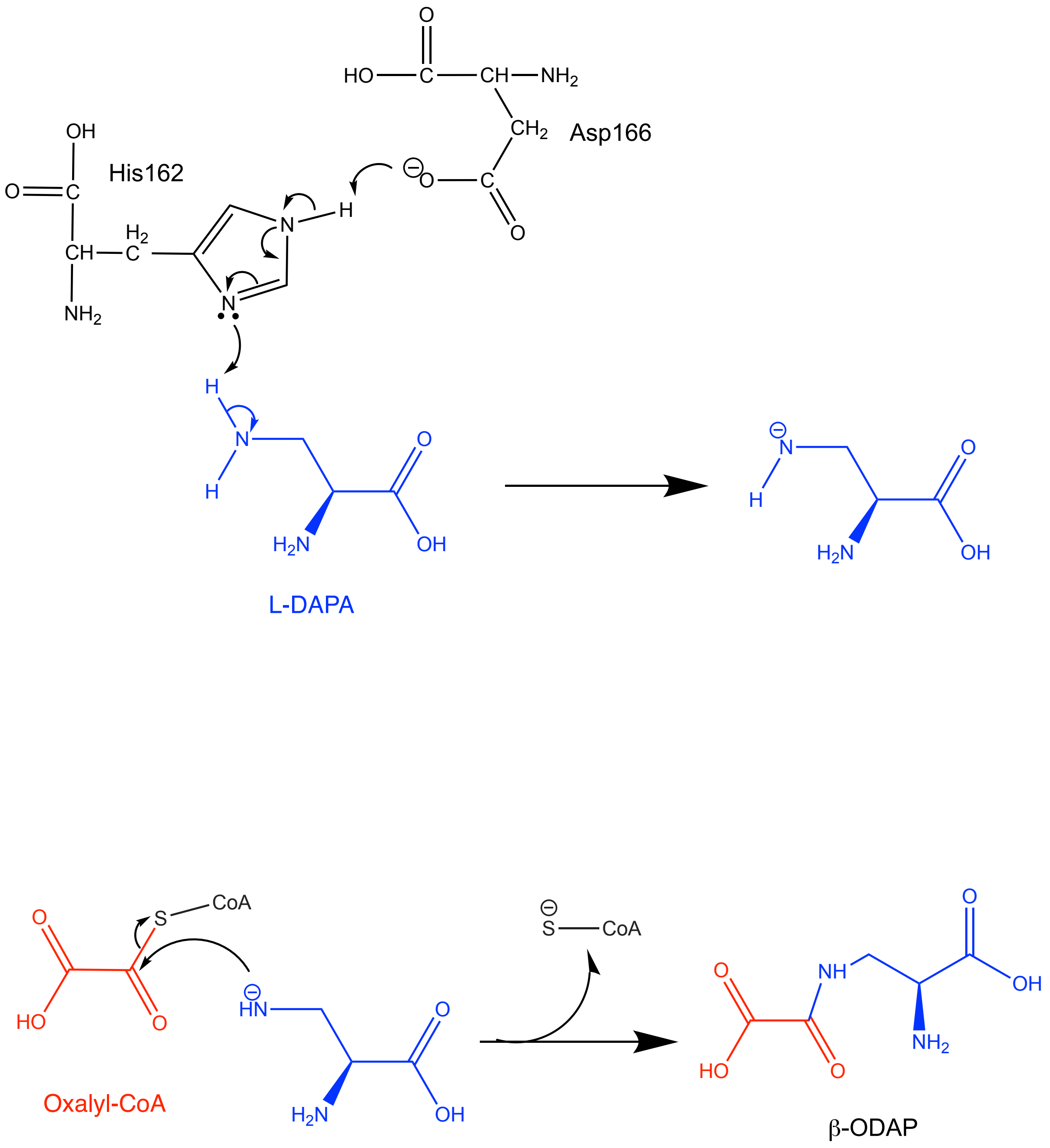


**Supplementary Figure S9. Proposed catalytic mechanism of BOS.** Asp166 increases the negative charge of His162 through deprotonation. His162 then acts as a general base that obstructs a proton from the β-amine group of L-DAPA (blue). The resulting nucleophile attacks the thio-ester carbonyl of oxalyl-CoA (red), forming β-ODAP and releasing CoA. This figure was created using ChemDraw version 16.0.1.4, PerkinElmer Informatics™.

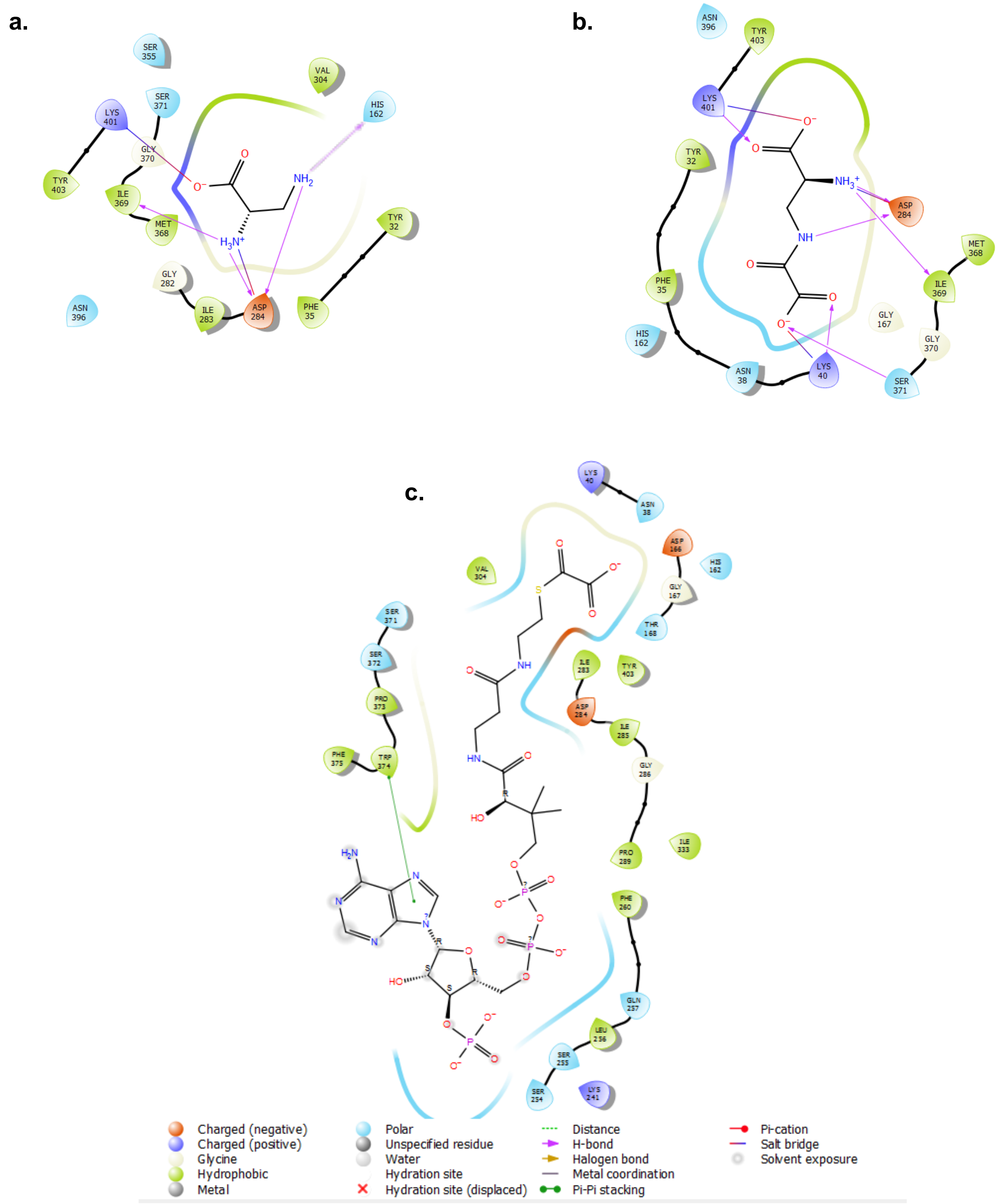


**Supplementary Figure S10. Two-dimensional ligand interaction maps of BOS with L-DAPA, oxalyl-CoA and β-ODAP.** Protein-ligand interactions were derived from individual docking studies of the ligand molecules to BOS. **a.** Binding interactions between L-DAPA and BOS. **b.** Binding interactions between β-ODAP and BOS. **c.** Binding interactions between oxalyl-CoA and BOS. The interactions were calculated and plotted as a 2D Ligand Interaction Diagram (Schrödinger)
